# Re-arrangements in the cytoplasmic distribution of small RNAs following the maternal-to-zygotic transition in *Drosophila* embryos

**DOI:** 10.1101/224626

**Authors:** Mehmet Ilyas Cosacak, Hatice Yiğit, Bünyamin Akgül

**Author notes:** **Correspondence:** Bünyamin Akgül, Ph.D., Molecular Immunology and Gene Regulation Laboratory, Izmir Institute of Technology, 35430, Gülbahçeköyü, Izmir, Turkey; **Email**.

## Abstract

Small RNAs are known to regulate gene expression during early development. However, the dynamics of interaction between small RNAs and polysomes during this process is largely unknown. 0-1h and 7-8h *Drosophila* embryos were fractionated on sucrose density gradients into four fractions based on A_254_ reading (1) translationally inactive messengerribonucleoprotein (mRNP); (2) 60S; (3) monosome; and (4) polysome. Comparative analysis of deep-sequencing reads from fractionated and un-fractionated 0-1h and 8-h embryos revealed development-specific co-sedimentation pattern of small RNAs with the cellular translation machinery. Although most miRNAs did not have a specific preference for any state of the translational machinery, we detected fraction-specific enrichment of some miRNAs such as miR-1-3p, -184-39, 5-5p and 263-5p. More interestingly, we observed dysregulation of a subset of miRNAs in fractionated embryos despite no measurable difference in their amount in unfractionated embryos. Transposon-derived endosiRNAs are over-expressed in 7-8h embryos and are associated mainly with the mRNP fraction. However, transposon-derived piRNAs, which are more abundant in 0-1h embryos, co-sediment primarily with the polysome fractions. These results suggest that there appears to be a complex interplay among the small RNAs with respect to their polysome-cosedimention pattern during early development in *Drosophila*.

## INTRODUCTION

In *Drosophila*, early developmental processes until the maternal-to-zygotic transition (MZT) require the function of nearly 7,000 maternally-loaded RNAs (1,2). The MZT transfers the developmental control to the zygotic genome and induces the degradation of maternal mRNAs. Accumulating evidence suggests that small non-coding RNAs (ncRNAs) play an essential role in regulating the MZT both in insects and mammals (3–7).

There are at least three types of well-documented small non-coding small RNAs (ncRNAs), microRNAs (miRNAs), endogenous silencing RNAs (endo-siRNAs) and PIWI-interacting RNAs (piRNAs) (8–10). miRNAs (miRNAs) are ubiquitously expressed small RNAs of ~22 nucleotide in length that regulate gene expression post-transcriptionally by decreasing RNA stability or repressing translation at the initiation, 60S joining or post-initiation steps (11–14). piRNAs appear to stabilize the germ cell genome and induce deadenylation of maternal mRNAs in the early *Drosophila* embryo(7,15). An abundance of sense and antisense endo-siRNAs are generated during transcription from different sites around the promoter and termination regions or from sense mRNAs and longer antisense transcripts (16). Endo-siRNAs can have various regulatory functions ranging from heterochromatin formation to post-transcriptional gene regulation (8).

There appears to be a complex interplay among the three major classes of small ncRNAs in regulating the MZT. The piRNA pathway has been reported to modulate maternal mRNA deadenylation and decay in the early *Drosophila* embryos (7,17). Both maternal and sperm-borne endo-siRNAs regulate fertilization and early developmental gene regulatory processes(18–20). Substantial changes have been observed in the miRNA expression profiles of embryos at different developmental stages in *Drosophila* (15,21,22) and targeted knockout mutations result in various phenotypes(23). The similarity in the miRNA expression pattern in mature oocyte and zygote suggests the presence of maternally inherited miRNAs(24). The significance of maternal miRNAs pre-MZT is largely unknown, except for miR-34, which is maternally inherited and important for neurogenesis in *Drosophila*(25). It is suggested that endo-siRNAs and miRNAs may be responsible for regulating early developmental processes before and after the MZT, respectively (26).

Coordinated regulation of mRNAs and miRNAs through the MZT stage is crucial for proper development. In *Drosophila*, most maternal RNA clearance is mediated by the RNA-binding protein Smaug(2). miR-309 plays a vital role in post-MZT elimination of maternal RNAs(5). Interestingly, maternal and zygotic miRNA expressions are coordinated as well. For example, Zelda, in conjunction with maternal morphogens and other zygotic transcription factors, trans-activates zygotic miRNA transcription (27) while Wispy is responsible for adenylation-dependent degradation of maternal miRNAs(28).

Despite a great progress in small RNA-mediated gene regulation, the control of small RNAs themselves has only been tackled recently(29,30). For instance, miRNA biogenesis can be regulated at multiple steps, including transcription, post-transcriptional processing, editing and intracytoplasmic localization. Apparently, there is a correlation between intracytoplasmic location of miRISC complexes and their function. Whereas, P-body-associated miRNAs may be involved in mRNA degradation (31) polysome-association of miRNAs may be correlated with miRNA-mediated translational regulation(32). Interestingly, polysome occupancy of miRNAs was shown to differ between human embryonic stem cells and foreskin fibroblast cells(33). Differential polysome occupancy, which is apparently influenced by the choice of seed not the abundance, is correlated with the target sequence. Even more interestingly, differential polysome association results in the formation of diverse miRNA effector complexes that are regulated by extracellular signalling (34).

We exploited the presence of extensive post-transcriptional gene regulatory networks that take place in *Drosophila* embryos to investigate intracellular dynamics of small RNAs pre- and post-MZT by deep-sequencing. Comparative analysis of fractionated and unfractionated 0-1- and 7-8h embryo cytosolic extracts showed that polysomal and non-polysomal fraction-associated small RNA profiles change dramatically following the MZT. Transposon-derived and intergenic-region-derived small RNAs are more abundant in 0-1h embryos while miRNAs are more abundant in 7-8h embryos in unfractionated embryos. miRNAs appear to interact with the translational machinery at all states, suggesting that each miRISC resides in distinct cytoplasmic reservoirs. Of the two types of transposon-derived small RNAs, siRNAs are expressed at 7-8h embryos and are primarily associated with polysomal complexes. piRNAs, on the other hand, are detected more abundantly at 0-1h embryos and primarily co-localize with non-polysomal complexes. Altogether these results suggest that embryos possess a different small RNA profile pre- and post-MZT and that each type of small RNAs possesses a different polysome association profile.

## MATERIAL AND METHODS

### Small RNA Deep-sequencing Data Analysis

Embryo collection, polysome profiling and small RNA deep-sequencing data have been previously described ((35), GEO accession number GSE35443). We used the small RNA-seq data to investigate the small RNA dynamics during early development in *Drosophila*. To this extent, the 36-bp multiplex sequences were split into their corresponding samples according to their barcodes. Only sequences without any ambiguity were used for further analyses.

After barcode selection, the 3′ adapter sequence was trimmed from the raw reads in 4 steps by using the 3′ adapter sequence 5′-ATCTCGTATGCCGTCTTCTGCTTGT-3′.(1) Trimming full adapter sequences identified inserts with a minimum length of 7 nt; (2) If no adapter sequences were found, the last base of the 3′ adapter was removed in a stepwise manner down to a minimum adapter size of 4 nt, which permit identifying inserts up to 8-28 nt; (3) Finally, the remaining reads were searched for adapter sequences with mismatches; (4) inserts with poly(A) and/or multiple “N”s were removed from the data. The resulting 15-29-bp inserts were collapsed into a fasta format and used for subsequent analyses.

For quality control analyses, nexalign program (36) was used to align all sequences to the genome (dmel-r5.39, flybase.org) and CCA-appended mature tRNAs (flybase.org). The sequences were categorized in an order as those with exact match (EMM), one (insertion or deletion or mismatch) (MINDEL), two (M2M) or three mismatches (M3M). The remaining sequences were grouped as unmapped (UNM). The number of mapping site in the genome and the mapping type were then appended to the header line of each unique read for further data analyses.

All known RNA sequences were downloaded from flybase (dmel-r5.39) except (i) hairpin and miRNAs sequences from mirbase (www.mirbase.org) (Release 17), (ii) rRNA sequences (5.8S, 18S and 28S) from NCBI (M21017.1), (iii) Repbase collection from Jurka et al. 2005(37), and (iv) piRNA cluster genomic coordinates from Brennecke et al. 2007(38). The piRNA clusters from sense strand were extracted by an in-house-algorithm from dmel-r5.39.

The 15-29-bp deep-sequencing reads were aligned to these known *Drosophila* RNAs in the following order: rRNA, miRNA hairpin, tRNA, miscRNA, ncRNA, transposon, transcript, intron, pseudogene and intergenic region. The initial alignment was carried out for exact matches followed by alignment with one, two or three mismatches. In order to analyse the miRNA cluster and Repbase collection, the transposon-mapped sequences were remapped for exact matches to the repbase collection and the intergenic-region- and transposon-mapped reads were realigned to the piRNA clusters.

To calculate miRNA expression levels, we used the exact-matched mature miRNAs sequences and hairpin sequences in miRBase (Release 17). The read per million (RPM) for each miRNA was calculated using the formula (fold change = ((7-8h_RPM + 10)/(0-1h_RPM + 10)) in which 10 reads were added to each read to eliminate overestimations in lower reads. The resulting miRNAs were clustered using Cluster 3.0 (39) and visualised by java Tree View(40). The transposon sequences from Flybase were used to identify the transposon-derived piRNA expression levels. We aligned the single-hit, transposon-derived 23-29-nt reads to the Repbase collection to obtain the log2 ratio of 0-1 and 7-8h RPM values. In order to validate that the 23-29-nt reads are indeed piRNAs, we searched for two features in these reads (i) a 10-nt 5′-5′ complementarity and (ii) a preference for U and A at the nucleotide positions 1 and 10, respectively. To analyse the piRNA clusters, the previously published piRNA cluster coordinates were used(38). The 21-nt reads were not used for piRNA analysis, as they represent siRNA population.

## RESULTS

Blastoderm cellularization, which takes place between 2.5 and 3h after fertilization in *Drosophila*, is the first developmental process that requires post-MZT zygotic transcription(2). Small RNAs play an important role in the regulation of the MZT to set the stage for post-MZT embryonic events(3,4,10). To gain insight into the expression levels of small RNAs during MZT, we analysed the small RNA-seq data that was previously deposited to GEO ((35), GSE35443). This data set includes small RNA-seq of 0-1- and 7-8h embryos. Half of these embryos were used to isolate total RNAs from unfractionated embryos (Figure 1A, UF) as a reference. The other halves were then fractionated, on sucrose density gradients (SDGs), into 4 major sub-fractions based on A254 absorbance: (1) messengerribonucleoproteins (mRNPs) devoid of rRNAs, (2) 60S, containing 28S rRNA, (2) monosome, and (4) polysome. Here we report the comparative analysis of dynamic changes that we have observed in the intracytoplasmic localization of all small RNAs during early development in *Drosophila*.

**Figure 1.**
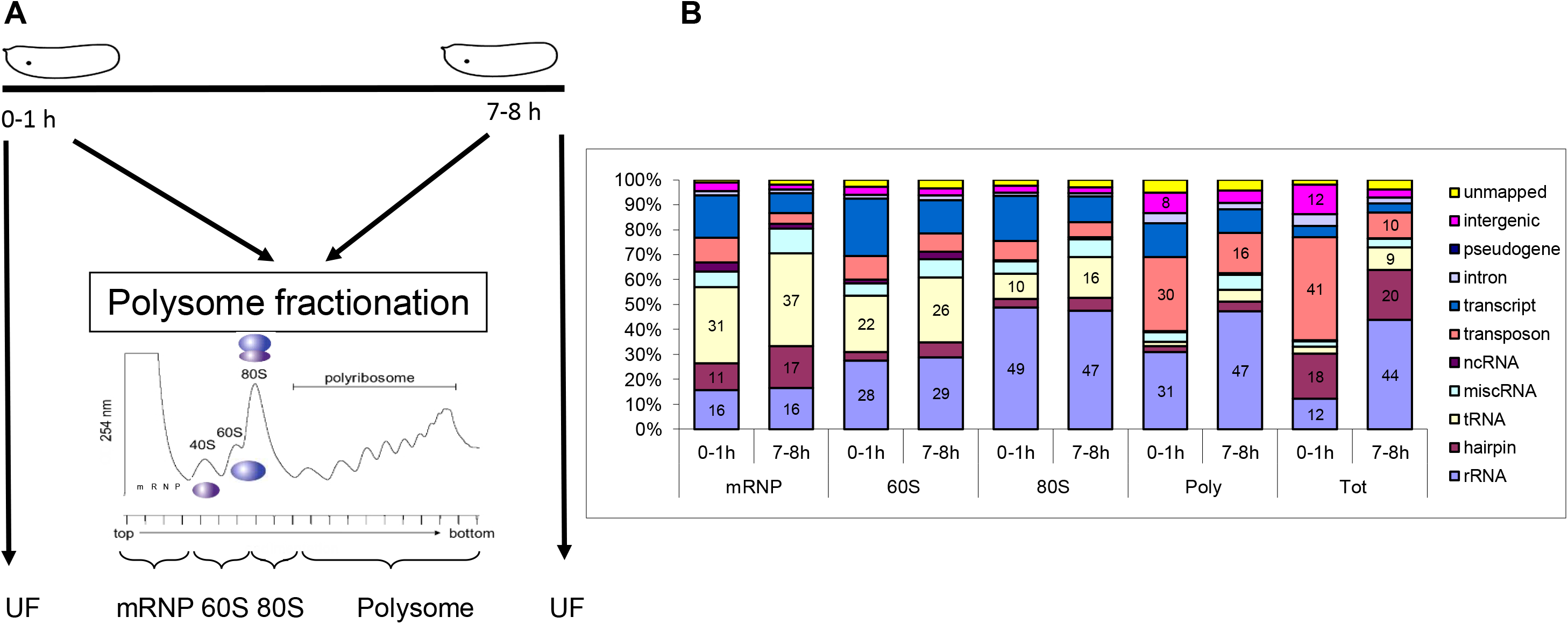
Alignment and categorization of small RNAs based on their origin. **(A)** Experimental design for total RNA extraction from un-fractionated (UF) and fractionated embryos has been published (35). **(B)** Percentage of small RNAs in un-fractionated and fractionated embryos. The sequences were aligned in an order from down to top as shown in the figure legend. The aligned sequences included not only the perfect matches but also indel and up to 3 mismatches.

One major concern in polysome profiling is the potential for a greater RNA degradation during sample processing prior to downstream events. Thus, we first checked the RNA quality of total RNAs and small RNAs using Agilent RNA 6000 Nano kit and Agilent Small RNA kit, respectively (Agilent, USA) (Figure S1). Total RNA capillary electrophoresis verified high quality of RNAs phenol-extracted from fractions(35). Interestingly, each fraction appears to possess a unique small RNA profile especially based on small RNAs that are smaller than tRNAs. When we calculated the frequency of deep-sequencing reads based on the read size, a great majority (88.8-99.5%) of adaptor-containing reads contained inserts of 15-29 bp in size, further verifying a high quality of input RNA. The low percentage of 0-14-bp inserts in fractionated RNAs, compared to that in unfractionated total RNAs, was consistent with the high quality of fractionated RNAs (Supporting Table 1). Interestingly, 1h total RNA contained the greatest percentage of 0-14bp inserts, congruous with the destabilization of maternal mRNAs (2). Of the 12,553,921 total reads, 71.95% of the fragments matched perfectly to the *Drosophila* genome with a number of unique sequences in each sample ranging from 71,830 to 757,803. Based on the relative length distribution, we observed enrichment in 22-28-bp RNAs (Figure S2).

### small RNA populations localize in distinct cytoplasmic reservoirs

We used the nexalign program to align sense or antisense sequences to the known RNAs(36). A pre-alignment to all known RNAs prompted us to align sequences in an order as described in the Methods section. We obtained a perfect match ratio of 71-82% whereas more than 90% of sequences mapped with exact or a single mismatch. Quite interestingly, each small RNA population appears to sediment with polysomal fractions to a different extent (Figure 1B). In the non-polysomal mRNP fraction, the majority of small RNAs contained tRNA-derived fragments (tRFs), with 31% and 37% in 0-1h and 7-8h, respectively. Based on northern blot results, we observed developmentally differentially expressed tRFs in accordance with the deep-sequencing data, which was published previously(35). In the polysomal fraction, the majority of small RNAs is derived from rRNAs and transposons. The pattern of enrichment with respect to the developmental stage (1h versus 8h) was similar in fractionated and unfractionated samples, indicating the potential biological significance rather than random degradation introduced during fractionation.

A relatively high percentage of rRNA-derived fragments in 80S and polysomal fractions of both 0-1- and 7-8h embryos is conceivable considering the abundance of ribosomes in these fractions (Figure 1B). We noticed a much higher rRNA-derived fragments in the total RNAs of 7-8h embryos compared to 0-1h embryos (Figure 1B) despite quite similar RNA quality based on Bioanalyzer capillary electrophoresis (Figure S1A). Interestingly, while the percentage of rRNA-derived fragments are similar in mRNP, 60S and 80S fractions, it is higher in polysomal fractions in 7-8h embryos. Normally, degraded fragments would be expected to enrich in mRNP fractions, however these small RNAs are mainly enriched in polysome, suggesting a biological degradation or processing from rRNAs rather than random degradation. Additionally, the transcript-derived fragments are more abundant in 0-1h embryos, which probably indicates either clearance of maternal mRNAs or biogenesis of novel zygotic small RNAs. We then focused primarily on miRNAs and transposon-derived small RNAs as they constitute the bulk of small RNAs in our samples.

### miRNAs interact with cellular translational machinery at all states

Although the temporal expression of miRNAs during early development is well documented(15), the extent to which each dys-regulated miRNA is associated with polysomes is unknown. To this end, we firstly aligned the RNA-seq reads to the miRNA hairpin with perfect matches. A total of 256 mature miRNAs perfectly matched with miRNA hairpins. Setting the minimal threshold at 50 RPM filtered out 162 miRNAs, resulting in a total of 94 mature miRNAs for further analyses.

To identify fraction-specific miRNAs, we selected ten most abundant hairpin sequences from each sample (10 hairpins from each sample of total, mRNP, 60S, monosome and polysome, yielding a total of 50 hairpins) and identified 18 common hairpin sequences that constituted 82.5-93.6% of all hairpin sequences with perfect matches in each sample (Table 1). Thus, these 18 miRNAs represent the most abundant miRNAs in each sample. We observed dynamic changes in the expression of many miRNAs in unfractionated total RNAs in agreement with the published results (Table 2, ref-15 (15)). Analysis of miRNA expression in fractionated samples yielded several interesting points (Table 1). Firstly, for the majority of miRNAs, the miRNA levels in fractionated samples were comparable to those in the unfractionated samples, probably indicating the non-selective distribution of these miRNAs throughout the cytoplasm. Secondly, although most miRNAs are distributed nearly evenly throughout the fractions, certain miRNAs were enriched in particular fractions. For instance, miR-5-5p is enriched in the mRNP fraction both in 0-1 and 7-8h embryos whereas miR-1-3p is enriched in the 80S fraction. Thirdly and more importantly, the extent to which a miRNA is associated with a particular fraction appears to be relatively similar for most miRNAs. However, for a few miRNAs, the degree of association is developmentally regulated. For instance, miR-9c-5p makes up of 11.5 and 4.1% of miRNAs in 0-1 and 7-8h embryos.

**Table 1.** The percentage of the most abundant 10 miRNAs. The sequences that perfectly matched to hairpin and mature miRNAs were used to calculate read frequency. The most abundant 10 miRNAs were selected from each sample and then pooled to generate a panel of 18 miRNAs that appear to be the most abundant miRNAs in 0-1- and 7-8h *Drosophila* embryos.

**Coşacak et al Table 1**
|  | hairpin |  |  |  |  |  |  |  |  |  | mature_3bp |  |  |  |  |  |  |  |  |  |
| --- | --- | --- | --- | --- | --- | --- | --- | --- | --- | --- | --- | --- | --- | --- | --- | --- | --- | --- | --- | --- |
|  | mRNP |  | 60S |  | 80S |  | Poly |  | Tot |  | mRNP |  | 60S |  | 80S |  | Poly |  | Tot |  |
|  | 0-1h | 7-8h | 0-1h | 7-8h | 0-1h | 7-8h | 0-1h | 7-8h | 0-1h | 7-8h | 0-1h | 7-8h | 0-1h | 7-8h | 0-1h | 7-8h | 0-1h | 7-8h | 0-1h | 7-8h |
| dme-bantam-3p | 0,3 | 3,4 | 36,3 | 27,3 | 35,9 | 26,4 | 2,9 | 1,0 | 1,5 | 30,3 | 0,3 | 3,3 | 35,7 | 26,2 | 35,7 | 26,1 | 2,6 | 0,9 | 1,5 | 30,1 |
| dme-miR-1-3p | 7,7 | 5,5 | 3,1 | 7,9 | 14,8 | 19,9 | 1,0 | 7,5 | 43,9 | 13,7 | 7,7 | 5,5 | 3,1 | 7,9 | 14,8 | 19,9 | 1,0 | 7,4 | 43,9 | 13,7 |
| dme-miR-184-3p | 17,5 | 9,6 | 7,6 | 4,6 | 10,4 | 6,1 | 27,0 | 25,9 | 11,1 | 3,6 | 17,5 | 9,6 | 7,6 | 4,6 | 10,4 | 6,1 | 27,0 | 25,9 | 11,1 | 3,6 |
| dme-miR-263a-5p | 18,0 | 24,7 | 1,5 | 4,2 | 3,4 | 5,1 | 33,1 | 22,2 | 3,5 | 9,0 | 18,0 | 24,7 | 1,5 | 4,2 | 3,4 | 5,1 | 33,1 | 22,1 | 3,5 | 9,0 |
| dme-miR-283-5p | 0,2 | 0,3 | 0,9 | 1,3 | 1,3 | 1,9 | 1,4 | 1,8 | 0,2 | 1,4 | 0,2 | 0,3 | 0,9 | 1,2 | 1,3 | 1,8 | 1,4 | 1,8 | 0,1 | 1,4 |
| dme-miR-286-3p | 2,0 | 10,8 | 7,1 | 18,1 | 7,5 | 18,9 | 6,8 | 10,3 | 2,2 | 11,6 | 2,0 | 10,8 | 7,1 | 18,1 | 7,5 | 18,9 | 6,8 | 10,3 | 2,2 | 11,5 |
| dme-miR-305-5p | 1,0 | 0,0 | 1,5 | 0,3 | 0,4 | 0,1 | 0,3 | 0,0 | 1,7 | 0,1 | 1,0 | 0,0 | 1,3 | 0,2 | 0,3 | 0,0 | 0,3 | 0,0 | 1,7 | 0,1 |
| dme-miR-306-5p | 2,7 | 0,0 | 0,1 | 0,0 | 0,1 | 0,0 | 0,1 | 0,0 | 2,4 | 0,1 | 2,7 | 0,0 | 0,1 | 0,0 | 0,1 | 0,0 | 0,1 | 0,0 | 2,4 | 0,1 |
| dme-miR-311-3p | 2,1 | 0,0 | 0,6 | 0,1 | 1,0 | 0,1 | 0,2 | 0,1 | 2,6 | 0,1 | 2,1 | 0,0 | 0,6 | 0,1 | 1,0 | 0,1 | 0,2 | 0,1 | 2,6 | 0,1 |
| dme-miR-315-5p | 0,2 | 0,8 | 0,6 | 0,8 | 0,3 | 0,5 | 1,4 | 0,8 | 0,2 | 2,0 | 0,2 | 0,8 | 0,6 | 0,8 | 0,3 | 0,5 | 1,4 | 0,8 | 0,2 | 2,0 |
| dme-miR-5-5p | 16,4 | 27,2 | 3,4 | 4,3 | 2,4 | 3,5 | 5,9 | 7,0 | 3,3 | 4,5 | 16,4 | 27,2 | 3,1 | 3,9 | 2,3 | 3,3 | 5,9 | 6,9 | 3,2 | 4,2 |
| dme-miR-7-5p | 0,1 | 1,5 | 0,5 | 1,2 | 0,6 | 1,6 | 1,1 | 1,4 | 0,3 | 1,0 | 0,1 | 1,5 | 0,5 | 1,2 | 0,6 | 1,6 | 1,1 | 1,4 | 0,3 | 1,0 |
| dme-miR-8-3p | 0,9 | 1,2 | 3,0 | 2,9 | 3,3 | 3,0 | 1,2 | 1,7 | 1,3 | 2,5 | 0,5 | 1,1 | 2,7 | 2,5 | 3,1 | 2,7 | 1,2 | 1,6 | 1,2 | 2,2 |
| dme-miR-92b-3p | 0,3 | 0,0 | 1,6 | 1,0 | 2,0 | 1,2 | 0,2 | 0,3 | 0,8 | 0,1 | 0,3 | 0,0 | 1,6 | 0,9 | 2,0 | 1,2 | 0,2 | 0,3 | 0,8 | 0,1 |
| dme-miR-958-3p | 0,0 | 0,1 | 4,2 | 5,1 | 0,5 | 0,6 | 0,1 | 0,2 | 0,0 | 0,8 | 0,0 | 0,1 | 4,1 | 5,0 | 0,5 | 0,6 | 0,0 | 0,1 | 0,0 | 0,8 |
| dme-miR-9a-5p | 6,0 | 5,3 | 1,6 | 3,5 | 1,7 | 2,9 | 1,7 | 8,6 | 4,1 | 6,4 | 6,0 | 5,3 | 1,6 | 3,4 | 1,7 | 2,9 | 1,7 | 8,6 | 4,1 | 6,4 |
| dme-miR-9c-5p | 11,5 | 4,1 | 8,8 | 2,4 | 6,0 | 1,8 | 9,1 | 4,4 | 9,4 | 2,6 | 11,5 | 4,1 | 8,8 | 2,4 | 6,0 | 1,8 | 9,0 | 4,4 | 9,4 | 2,5 |
| dme-miR-iab-4-5p | 1,2 | 0,1 | 0,1 | 0,1 | 0,0 | 0,0 | 0,1 | 0,0 | 0,5 | 0,1 | 1,6 | 0,1 | 0,0 | 0,1 | 0,0 | 0,0 | 0,2 | 0,1 | 0,7 | 0,1 |

**Table 2.** Categorization of miRNAs based on their association with translational machinery. Based on fold of induction between 0-1- and 7-8h embryos, miRNAs were divided into 4 major miRNA groups (Group 1-overexpressed in 7-8h unfractionated embryos; Group 2-equally expressed in unfractionated embryos; Group 3-downregulated in 7-8h unfractionated embryos; Group 4; Others). The fold changes in red, green and light blue indicate increase, decrease and no significant change in 7-8h embryos, respectively.

**Coşacak et al Table 2**
| G | miRNA | Tot | mRNP | 60S | 80S | Poly | G | miRNA | Tot | mRNP | 60S | 80S | Poly |
| --- | --- | --- | --- | --- | --- | --- | --- | --- | --- | --- | --- | --- | --- |
| 1 | dme-miR-286-3p | 2,6 | 3,1 | 2,2 | 2,1 | 1,6 | 2 | dme-miR-11-3p | 0,8 | 1,3 | 0,8 | 0,5 | 2,4 |
| 1 | dme-miR-283-5p | 3,6 | 1,5 | 1,3 | 1,3 | 1,3 | 2 | dme-miR-9b-5p | 0,4 | 0,8 | -0,4 | -0,2 | 1,7 |
| 1 | dme-miR-7-5p | 2,0 | 4,5 | 2,0 | 2,1 | 1,3 | 2 | dme-miR-1012-3p | -0,8 | -0,8 | -0,5 | -1,0 | 1,4 |
| 1 | dme-miR-957-3p | 2,6 | 1,3 | 1,2 | 1,2 | 0,8 | 2 | dme-miR-1010-3p | 0,2 | -0,6 | 0,4 | 0,5 | 1,4 |
| 1 | dme-miR-263a-5p | 1,6 | 1,1 | 2,3 | 1,3 | 0,4 | 2 | dme-miR-5-5p | 0,6 | 1,4 | 1,2 | 1,3 | 1,2 |
| 1 | dme-miR-315-5p | 3,9 | 2,6 | 1,3 | 1,3 | 0,2 | 2 | dme-miR-12-5p | 0,3 | 0,4 | 0,6 | 1,2 | 1,1 |
| 1 | dme-miR-314-3p | 5,6 | 2,1 | 1,3 | 1,0 | 0,0 | 2 | dme-miR-13a-3p | 0,2 | 0,6 | 1,3 | 0,2 | 0,0 |
| 1 | dme-bantam-5p | 2,6 | 1,7 | 1,5 | 1,2 | -0,8 | 2 | dme-miR-31a-3p | 0,2 | 0,0 | 1,0 | 0,7 | -0,3 |
| 1 | dme-miR-958-3p | 7,0 | 3,6 | 1,1 | 0,8 | 1,8 | 2 | dme-miR-124-3p | 0,0 | -0,3 | 1,2 | 0,2 | -0,3 |
| 1 | dme-miR-956-3p | 3,3 | 1,2 | 1,0 | 0,6 | 0,7 | 2 | dme-miR-965-5p | 0,9 | -0,4 | 0,7 | 0,8 | 0,8 |
| 1 | dme-miR-998-3p | 1,3 | 1,2 | 1,5 | 0,5 | 0,4 | 2 | dme-miR-1003-3p | -0,8 | -0,5 | 0,1 | 0,0 | 0,0 |
| 1 | dme-miR-2c-5p | 1,4 | 1,2 | 1,6 | 0,2 | 0,2 | 2 | dme-miR-190-5p | 0,4 | 0,1 | 0,0 | 0,1 | -0,7 |
| 1 | dme-miR-304-5p | 6,0 | 1,5 | 0,9 | 0,5 | 0,1 | 2 | dme-miR-282-3p | -0,1 | -1,9 | -1,7 | -0,7 | -0,7 |
| 1 | dme-miR-8-3p | 1,1 | 1,8 | 0,8 | 0,5 | 1,3 | 2 | dme-miR-2a-3p | 0,6 | 0,2 | 0,8 | 0,5 | -1,5 |
| 1 | dme-miR-983-5p | 2,0 | 1,2 | 0,7 | 0,3 | 0,0 | 2 | dme-miR-33-5p | -0,3 | -2,4 | -1,6 | -1,1 | -1,9 |
| 1 | dme-bantam-3p | 4,6 | 4,3 | 0,4 | 0,3 | -0,5 | 2 | dme-miR-281-2-5p | -0,5 | -1,3 | 0,3 | -0,1 | 0,6 |
| 1 | dme-miR-2b-3p | 1,6 | 1,2 | 0,4 | 0,8 | -1,1 | 2 | dme-miR-2b-2-5p | -0,8 | -1,6 | 0,5 | 1,0 | 0,5 |
| 1 | dme-miR-276a-3p | 3,3 | 0,9 | 1,4 | 1,0 | 0,8 | 2 | dme-miR-375-3p | 0,1 | -2,3 | 0,8 | -0,2 | 0,1 |
| 1 | dme-miR-5-3p | 2,7 | 0,8 | 1,7 | 1,0 | 2,2 | 3 | dme-miR-3-5p | -2,8 | 0,0 | -0,1 | -0,1 | 0,0 |
| 1 | dme-miR-263b-5p | 1,5 | 0,7 | 1,9 | 1,2 | 1,5 | 3 | dme-miR-184-3p | -1,4 | -0,2 | 0,1 | 0,0 | 0,9 |
| 1 | dme-miR-137-3p | 1,3 | -0,6 | 1,6 | 1,3 | 1,4 | 3 | dme-miR-312-5p | -3,8 | -0,2 | -1,1 | -0,6 | -0,6 |
| 1 | dme-miR-1002-5p | 3,1 | -0,2 | 1,2 | 1,9 | 1,4 | 3 | dme-miR-6-1-5p | -4,2 | -0,3 | -0,7 | -0,8 | 0,0 |
| 1 | dme-miR-10-5p | 1,9 | 0,2 | 1,2 | 0,7 | 2,2 | 3 | dme-miR-3-3p | -1,8 | -0,4 | 0,4 | 0,5 | 0,7 |
| 1 | dme-miR-252-5p | 1,5 | 0,8 | 1,7 | 0,7 | 0,5 | 3 | dme-miR-999-3p | -1,7 | -0,4 | -0,2 | -0,1 | 0,6 |
| 1 | dme-miR-316-5p | 3,4 | 0,0 | 1,3 | 0,9 | -0,1 | 3 | dme-miR-305-3p | -1,0 | -0,7 | -0,2 | -0,6 | 0,0 |
| 1 | dme-miR-1006-3p | 2,5 | 0,5 | 1,1 | 0,5 | 0,0 | 3 | dme-miR-9c-5p | -1,7 | -0,8 | -1,0 | -1,0 | 0,0 |
| 1 | dme-miR-968-5p | 4,2 | 0,6 | 1,1 | 0,3 | 0,0 | 3 | dme-miR-13b-2-5p | -1,4 | -0,8 | 0,7 | 0,6 | 0,1 |
| 1 | dme-miR-1000-5p | 1,9 | 0,1 | 1,1 | 0,6 | 0,7 | 3 | dme-miR-9b-3p | -2,5 | -1,1 | 0,0 | -0,1 | -0,3 |
| 1 | dme-miR-987-5p | 2,1 | 0,7 | 0,2 | 0,1 | 1,1 | 3 | dme-miR-79-3p | -2,3 | -1,1 | -0,6 | 0,0 | 0,7 |
| 1 | dme-miR-31a-5p | 1,3 | 0,9 | 0,5 | 0,1 | 0,8 | 3 | dme-miR-92a-3p | -2,3 | -1,3 | 0,3 | -0,3 | 1,4 |
| 1 | dme-miR-13b-3p | 1,1 | 0,1 | 0,9 | 0,7 | 0,5 | 3 | dme-miR-124-5p | -1,5 | -1,3 | 0,5 | 0,5 | 0,4 |
| 1 | dme-miR-927-3p | 1,8 | 0,6 | 0,6 | 0,6 | 0,5 | 3 | dme-miR-1002-3p | -1,6 | -1,6 | 0,3 | 0,2 | 0,0 |
| 1 | dme-miR-996-3p | 1,0 | -0,2 | -0,2 | 0,2 | 0,1 | 3 | dme-miR-308-3p | -1,4 | -1,7 | -0,8 | -0,1 | -0,5 |
| 1 | dme-miR-284-5p | 2,4 | 0,4 | 0,6 | 0,1 | 0,0 | 3 | dme-miR-310-3p | -3,0 | -1,7 | -1,4 | -1,7 | -0,3 |
| 1 | dme-miR-927-5p | 2,1 | -0,6 | 0,5 | 0,4 | 0,0 | 3 | dme-miR-92b-3p | -2,5 | -1,9 | 0,1 | 0,0 | 1,2 |
| 1 | dme-miR-87-3p | 2,6 | -0,3 | 0,7 | 0,0 | -0,1 | 3 | dme-miR-968-3p | -2,1 | -2,2 | 0,3 | 0,5 | 0,7 |
| 1 | dme-miR-281-3p | 1,6 | -0,2 | -0,2 | -0,7 | -0,1 | 3 | dme-miR-995-3p | -1,4 | -2,4 | -0,1 | -0,4 | 0,4 |
| 1 | dme-miR-1008-3p | 1,4 | -0,2 | 0,5 | 0,3 | -0,5 | 3 | dme-miR-312-3p | -5,2 | -2,5 | -1,2 | -0,9 | -0,2 |
| 1 | dme-miR-8-5p | 1,1 | -1,6 | 1,0 | 1,0 | 2,6 | 3 | dme-miR-279-3p | -3,9 | -2,6 | 0,3 | 0,0 | 0,8 |
| 1 | dme-miR-10-3p | 2,2 | -1,0 | 0,6 | 1,2 | 2,0 | 3 | dme-miR-6-2-5p | -4,9 | -2,7 | -1,0 | -0,1 | 0,4 |
| 1 | dme-miR-14-3p | 1,0 | -1,4 | 1,5 | -0,3 | 1,1 | 3 | dme-miR-iab-8-3p | -2,3 | -3,2 | 0,8 | 0,5 | -0,3 |
| 4 | dme-miR-1-3p | -1,5 | 0,2 | 2,2 | 1,2 | 3,8 | 3 | dme-miR-iab-4-5p | -2,3 | -3,3 | 1,0 | 0,7 | -0,3 |
| 4 | dme-miR-988-3p | -1,0 | -0,1 | 1,4 | 1,2 | 0,0 | 3 | dme-miR-275-3p | -2,9 | -3,6 | -1,2 | -1,4 | 0,2 |
| 4 | dme-miR-4-3p | -1,1 | -0,5 | 2,2 | 0,7 | 1,4 | 3 | dme-miR-282-5p | -2,5 | -3,7 | -1,7 | -1,2 | -1,4 |
| 4 | dme-miR-31b-5p | -1,2 | -0,9 | 0,1 | -0,9 | 1,7 | 3 | dme-miR-311-3p | -5,2 | -4,8 | -1,2 | -2,0 | -0,2 |
| 4 | dme-miR-981-3p | 1,0 | -1,9 | -0,1 | 0,6 | 0,4 | 3 | dme-miR-305-5p | -4,0 | -4,9 | -1,8 | -1,8 | -1,3 |
| 2 | dme-miR-9a-5p | 0,8 | 0,5 | 1,9 | 1,5 | 3,3 | 3 | dme-miR-306-5p | -5,1 | -5,2 | -0,6 | -0,8 | -1,1 |

It is well known that maternal miRNAs are degraded after the MZT is completed and zygotic miRNA transcription modulates zygotic gene regulatory networks (28). It is unknown however whether the polysome status of zygotic miRNAs follow that of maternal miRNAs. There are potentially two possibilities (1) if the target sequence determines the mode of action of miRNAs, we would expect differential polysome association pre- and post-MZT, assuming that the targets of miRNAs pre- and post-MZT stage are different; (2) if the mode of action of miRNAs is independent from the target sequences, then we would expect to see a similar polysome-association pattern. To differentiate between these two hypotheses, we first calculated the miRNA expression ratios (0-1h/7-8h) in the unfractionated and fractionated samples. We then checked how the ratio in the unfractionated samples is reflected upon that in the fractionated samples. We assumed that if the cytoplasmic fate of a miRNA does not change pre- and post-MZT, the ratios obtained from the unfractionated and fractionated samples should be similar between 0-1- and 7-8-h embryos. In contrast, if there are any changes in the localization of miRNAs, the difference in the expression level obtained from the unfractionated samples should manifest in a particular sub-cellular fraction. This approach resulted in identification of 4 different miRNA groups with each having unique expression behaviour.

The first group includes 41 miRNAs that are over-expressed in 7-8h embryos (Table 2, G1; Figure 2). While 17 miRNAs co-sediment with the complexes in the mRNP fraction, some co-sediment with the 60S fraction. Interestingly, for 9 miRNAs, we did not see a proportional increase in their expression in fractionated RNAs although their expression was increased in un-fractionated embryos, suggesting the involvement of nuclear retention or other unknown mechanisms. The relative amount of miRNAs classified in the second group does not change in 0-1h and 7-8h un-fractionated embryos, indicating a similar transcriptional activity and/or miRNA stability. However, we detected dynamics changes in their subcellular locations following the MZT (Table 2, G2; Figure 2). For instance, despite no difference in the total miR-9a-5p amount in un-fractionated embryos, this specific miRNA becomes more polysome associated in 7-8h embryos (Log2 fold = 3.3). The third group includes 29 miRNAs, whose expression decreases in 7-8h embryos (Table 2, G3; Figure 2). Interestingly, the decrease in the miRNA expression is not distributed evenly throughout the 4 fractions, suggesting a preference for a specific fraction. For instance, 20 out of these 29 miRNAs sediment specifically with non-polysomal fractions, particularly in the mRNP fraction. This indicates the majority of small RNAs acts at mRNP complexes in early embryo compared with 7-8h. The fourth group includes the miRNAs whose relative abundance in the un-fractionaed and fractionated embryos are different. These miRNAs behave similarly to those in the second group in that the transcriptional output from these miRNAs do not lead to proportional changes in their cytoplasmic location, suggesting differential localization pathways pre- and post-MZT (Table 2, G4; Figure 2).

**Figure 2.**
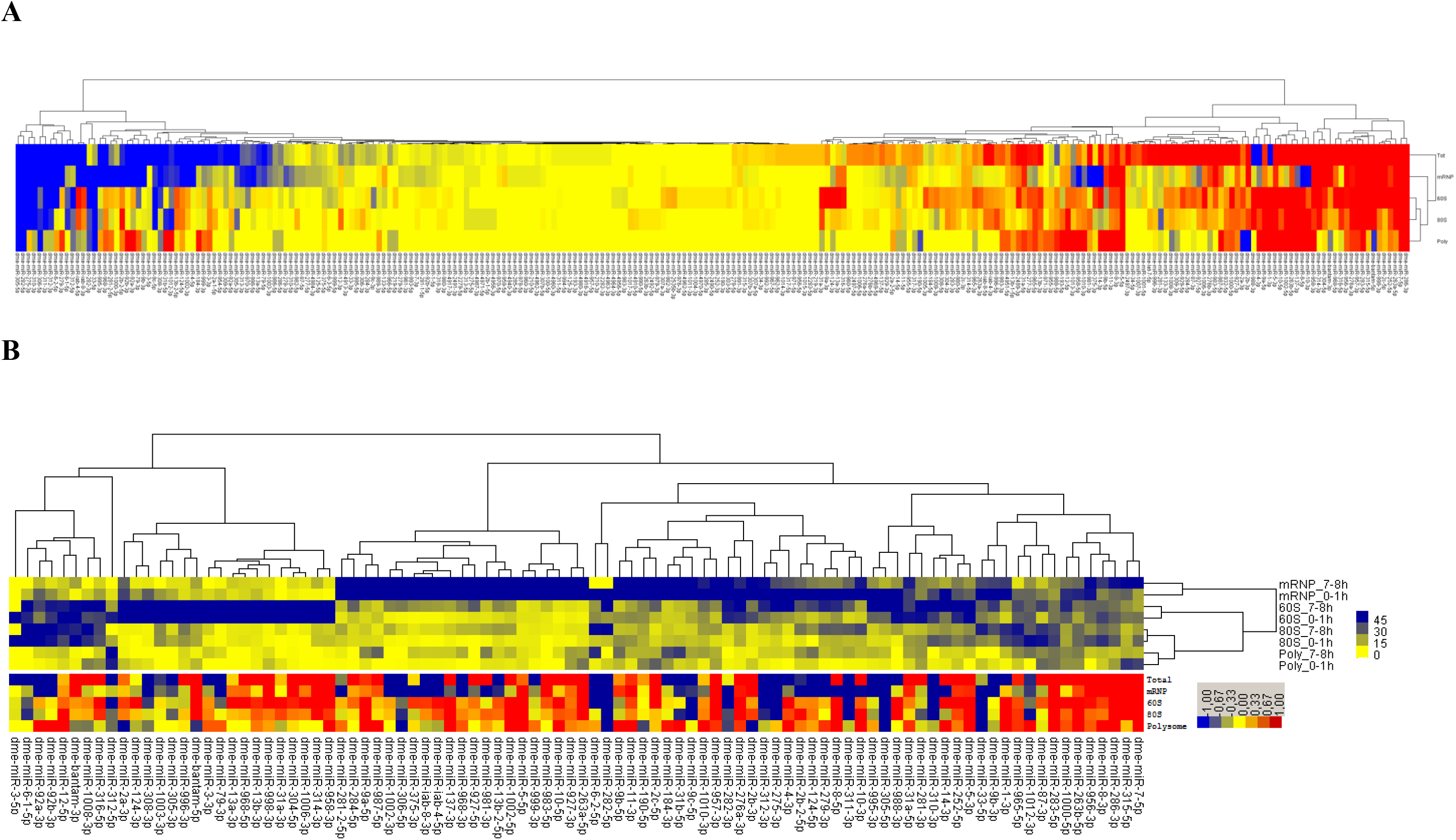
The Clustering of miRNA expression level. RNA-seq reads were first aligned to mature miRNA sequences with perfect matches, which yielded a total of 256 mature miRNAs. Setting the minimal threshold at 50 RPM filtered out 162 miRNAs, resulting in a total of 94 mature miRNAs. The log2 fold changes of these 94 miRNAs were then clustered using Gene Cluster 3.0(39) and visualized by Java Tree View (40) (A).Relative distribution of miRNA in fractions. In order calculate the relative abundance of miRNAs in each fraction, we calculated the percentage of each mature miRNA-mapped reads in each fraction hairpin mapped reads (B).

### miRNA Cluster Behaviours and miRNA Editing

Some miRNAs are known to be transcribed as members of gene clusters(41). Expression studies revealed that miRNA clusters are co-expressed (42) and cluster members are coordinated during target regulation(43). Assuming such coordination during target RNA regulation, we hypothesized that the cytoplasmic fate of the members should be similar. Thus, we compared the extent to which each cluster member is associated with four different fractions in our experimental design, each representing a different translational state of the cell. Our cluster analysis showed that miRNA cluster members behave similarly with respect to their cytoplasmic localization.

We also checked the frequency of post-transcriptional miRNA editing events likely to occur during the early development in *Drosophila* as miRNA editing is commonly used in eukaryotes to modulate the targets of miRNAs (44). We first aligned our sequences to the known miRNA sequences and looked for the sequences that align to the known sequences with a single mismatch. Based on this approach, we identified one candidate editing event (dme-miR-986, C→T at 11^th^ position). The PCR-amplification and sequencing of dme-miR-986 from P2 strain embryos and S2 cells showed that this particular difference in the sequence stems from an SNP not an editing process (data not shown).

### Transposon-derived siRNAs and piRNAs interact with different complexes

Some piRNAs are maternally deposited into the oocyte and they might be involved in the degradation of maternal mRNAs from the embryo(7). Also, MILI/MIWI is associated with polysomal fractions(45). However intra-cytoplasmic distribution and re-arrangements, if any, of piRNAs during early development are unknown. We used the Repbase collection to calculate the small RNAs generated from transposons. The expression of transposon-derived small RNAs decreased towards 7-8h, suggesting the significance of these small RNAs probably before the MZT. To ensure that we selected the transposon-derived piRNAs, we looked for two main features associated with piRNAs. The first feature is the 10-nucleotide complementarity at their 5′ ends as generated by the ping pong cycle (Figure 3A). The second feature in *Drosophila* is the presence of a U nucleotide at the 1st position and an A nucleotide at the 10th position (Figure 3B; (46)). Furthermore, we only used the reads in length of 23-29 bp to select the sequences that match the aforementioned two criteria.

**Figure 3.**
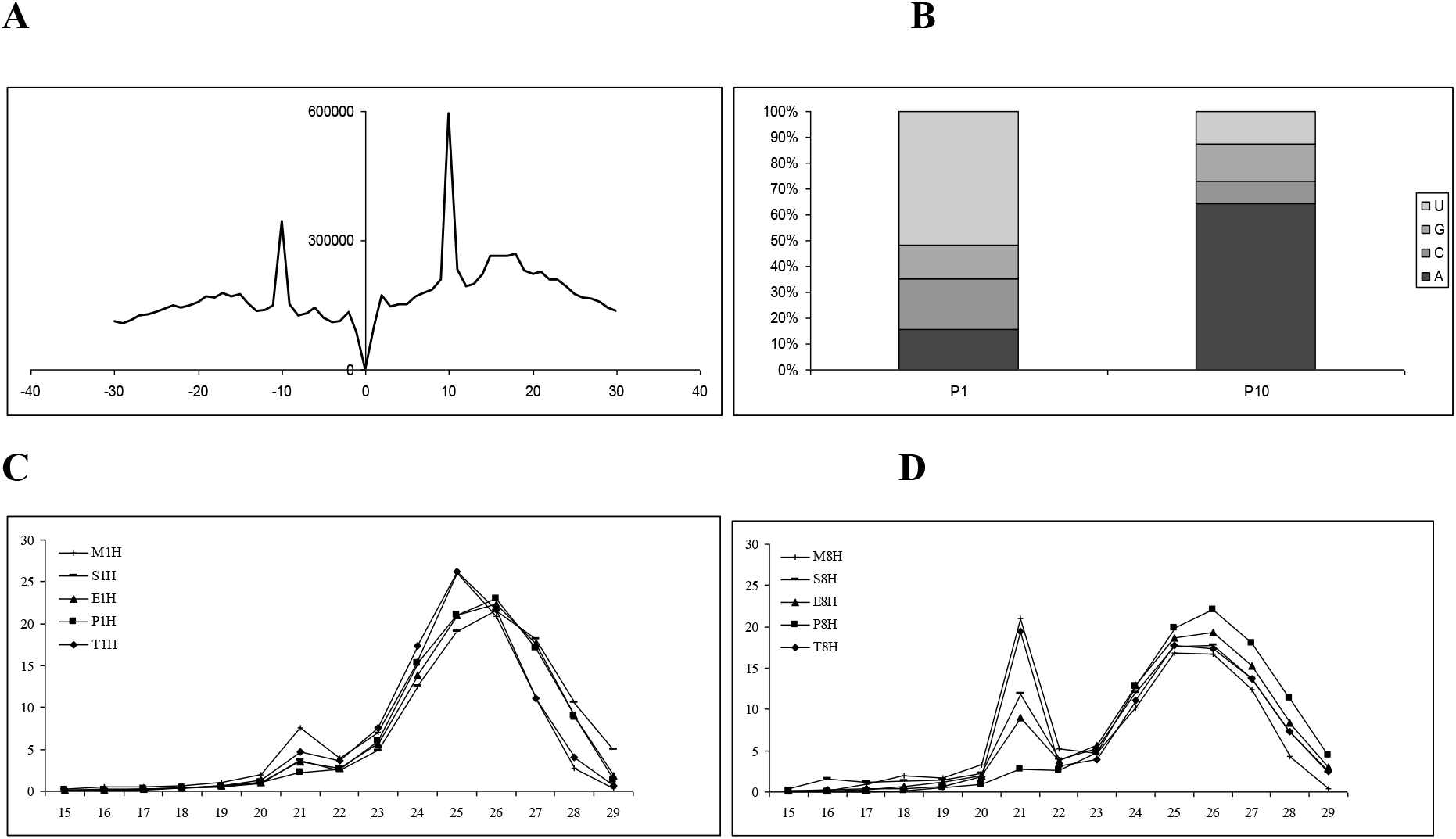
Analysis of transposon-derived transcripts. **(A)** The 5′-5′ complementarity among the transposon matched small RNAs. The complementarity of reads derived from transposon derived repeats were calculated by looking at 5’-to-5’ complementarity. The frequency at each length of complementation was plotted. The highest complementation occurs in the length of 10 nt. **(B)** The nucleotide bias at position 1 and 10 of the transposon-derived small RNAs. The small RNAs that have a 10-nt 5’-to-5’ complementation from Panel A were selected for calculating nucleotide frequency at each position. P1, Nucleotide position 1; P10, nucleotide position 10. We observed an enrichment for the U and A nucleotides at positions 1 and 10, respectively. The nucleotide distribution of transposon-derived small RNAs in 0-1h **(C)** and 7-8h embryos **(D)**.

We divided transposon-derived small RNA transcripts into groups based on their sizes: transposon-derived piRNAs of 23-29 nt (47) and transposon-derived siRNAs of 21 nt (47). To globally compare the polysome association of transposon-derived piRNAs to that of transposon-derived siRNAs in embryos, we calculated the read frequency of transposon-derived small RNAs. We noticed a much higher peak at the 21 bp reads in 7-8h embryo total RNAs (Figure 3D) compared with 0-1h (Figure 3C). Moreover, the 21-bp read frequency is higher in mRNP-associated complexes while decreasing towards polysomal fractions. This data suggest that transposon-derived siRNA expression is more abundant in later developmental stages although piRNA expression appears to be slightly higher in 0-1h embryos. Additionally, the intracellular localization of transposon-derived siRNAs appears to be different from that of transposon-derived piRNAs (Figure 3C–D). Although transposon-derived piRNAs are distributed throughout four fractions nearly equally, transposon-derived siRNAs are predominantly associated with the mRNP fraction. We then checked the frequency of reads from the 42AB cluster as it is one of the best characterized piRNA clusters in *Drosophila*(48). This analysis showed, in parallel to previous findings, production of transcripts from both strands: Interestingly, there appears to be a strand-bias in polysome-associated transcripts especially in 0-1h embryos (Figure 4B).

**Figure 4.**
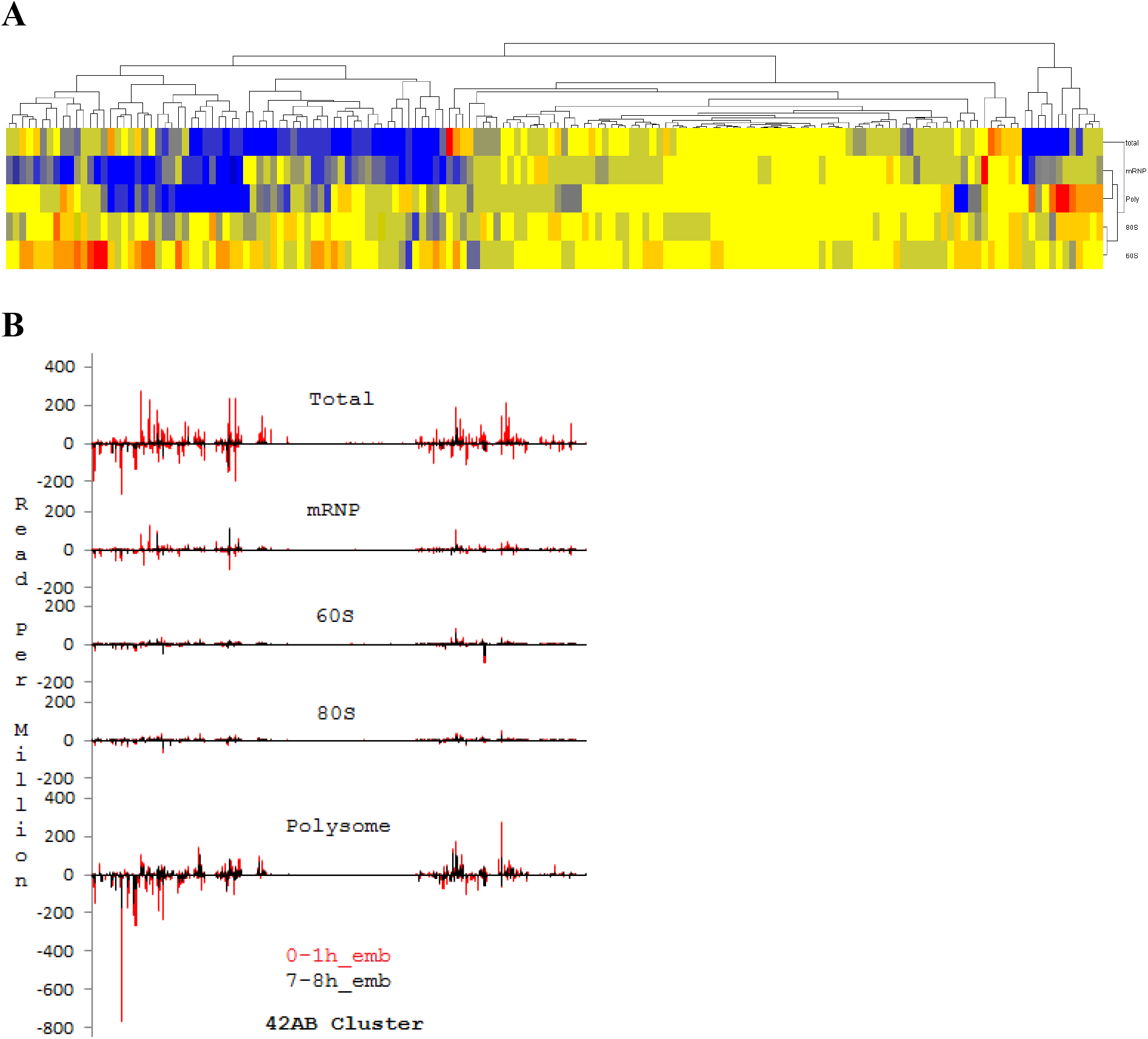
The Clustering of transposon-derived transcripts. RNA-seq reads were first aligned to the Repbase collection and uniquely mapped reads were used to calculate the transcript expression levels. Then the log2 fold change was calculated and clustered as in Figure-2A.

To substantiate our observation that transposon-derived piRNAs are highly expressed earlier compared to the siRNAs, we collected previously published deep-sequencing data from different developmental stages of *Drosophila melanogaster*(15,38,49). We used the same strategy as in Figure 4 to trace the temporal expression of siRNAs and piRNAs. Our analysis showed that the siRNA expression is relatively low in early developmental stages but increases significantly during later developmental stages (Figure 5A). Using the same data sets, we also checked the relative abundance of *Drosophila* small RNAs in various developmental stages. This analysis validated the notion that the transposon-derived small RNA abundance drops in later developmental stages while miRNA expression level increases (Figure 5B). Moreover, the 21-nt siRNAs derived from transposon increase while the piRNA levels (23-29nt) decrease.

**Figure 5.**
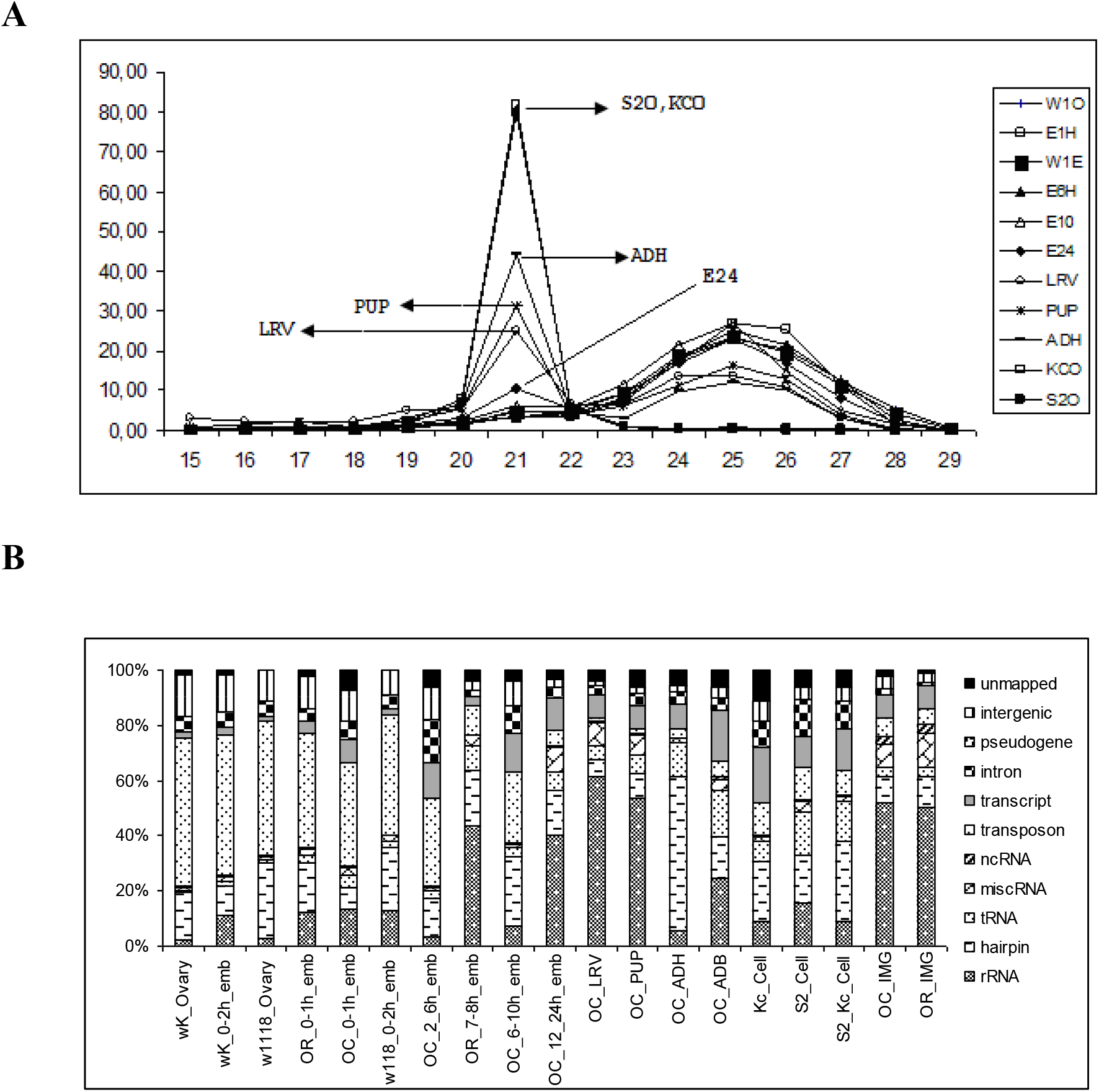
Analysis of small RNA abundance in various developmental stages. **(A)** The percentage of transposon-derived small RNAs in different developmental stages of *Drosophila*. The length distribution of transposon matched small RNAs in percentage during developmental stages of *Drosophila*. **(B)** The small RNAs aligned to know RNAs as in Figure-1B for all datasets downloaded from GEO. The strain names are wK: wild type caugth, OR: laboratory wild type, W1118: white eye, OC: Oregon Cansas, Ovary: oocyte, emb: embryo, Kc and S2: Drosphila cell lines, ADH: Adult Head, ADB: Adult Body, LRV: Larva, PUP: Pupa, IMG: Imaginal Discs. (These data sets were collected from previously published articles and raw sequences downloaded from GEO according to the GEO Accession Numbers in these articles; (15,38,49).

## DISCUSSION

We present here an in-depth analysis of cytoplasmic distribution of small RNAs in early development with respect to their association with the cellular translational machinery. The comparative analysis of the most abundant small RNAs reveals that the spatial location of each type of small RNA is quite different. Interestingly, miRNAs appear to utilize numerous molecular mechanisms as they interact with the translational machinery at all states. piRNAs, which are expressed more abundantly at the pre-MZT stage, are associated with polysomal complexes. In contrast to piRNAs, transposon-derived siRNAs are more abundantly expressed at the post-MZT stage and are primarily found in the mRNP fraction.

During the maternal-to-zygotic transition two crucial events take place: (i) clearance of maternal mRNAs and (ii) transcriptional activation of the zygotic genome. The clearance of maternal mRNAs is orchestrated by numerous RNA-Binding proteins, including SMAUG (1,2) and small RNAs such as miRNAs, siRNAs and piRNAs (4,7,50,51). Differential expression and specific functions of miRNAs in early development were previously reported (15,22) and our findings are in agreement with these results. For instance, miR-8 and 10 are up-regulated in 7-8h embryos in total RNAs while miR-310 and 311 are down-regulated.

Although the differential expression of miRNAs during the MZT stage is well documented, the polysome profile of miRNAs during this switch is unknown. Our findings suggest that the majority of miRNAs appear to be associated with distinct fractions and probably with distinct complexes as a result (e.g., mRNP or polysomal complexes) (Table 2). For instance, miR-263-5p, 5-5p and 9c-5p are primarily found in the mRNP fraction whereas miR-184-3p is mainly part of polysomal fractions. Bantam-3p and miR-1-3p are primarily enriched in 60S and 80S fractions, respectively (Table 2). Although we cannot attribute any mechanistic insight into the fraction-specific functions of miRNAs yet, it might be noteworthy to speculate that this specific localization might be associated with the mode of actions of miRNAs. For instance, since mRNP complexes are known to include translationally inactive mRNAs (52), the miRNAs in this fraction might be involved in sequestration of mRNAs (or storage) away from the translational machinery. The miRNAs in the 60S and 80S fractions are likely to interfere with the translation initiation while the polysome-associated miRNAs probably interfere with translation elongation.

One other interesting observation from our findings is that the change in the amount of miRNAs in total RNAs (transcriptional or post-transcriptional increases/decreases) is not distributed equally throughout each fraction in the embryos. Some of differentially expressed miRNAs are directed towards the translationally inactive mRNP complexes whereas some others are destined for the translationally active polysomal complexes (Table 2). We interpret this observation to mean that miRNAs have a pre-defined cytoplasmic fate following their nucleo-cytoplasmic transport. Based on the differential cytoplasmic fate of miRNAs, it is appealing to propose that the composition of each miRISC complex might be quite versatile. It requires further investigation, however, to unravel what governs the cytoplasmic fate of miRISC complexes. More interestingly, for a few miRNAs (Table 2, miR-1-3p, miR-2b-3p, miR-92a-3p, miR-92b-39 and miR-1012-3p) miRISC complexes appear to switch from one translational state to another. For example, mRNP-fraction-associated miR-1-3p switches to polysomal fractions following the MZT. We cannot, however, conclusively claim that miRNAs switch their intracellular location as some of these post-MZT miRNAs could be zygotically transcribed miRNAs. In this situation, there should be additional factor(s), e.g., miRNA-binding proteins that specify the polysome status of zygotically transcribed miRNAs.

In either case, our data suggests that the activity of miRNP complexes may be spatially modulated through the differential cytoplasmic localization of these complexes pre- and post-MZT. More direct evidence is required, however, to demonstrate whether the eukaryotic cells utilize intracellular re-localization of miRNPs as a means to modulate miRNA function and target mRNA(s) as a result.

We also detected cytoplasmically localized transposon-derived small RNAs, which are classified as siRNAs (21 nt) or piRNAs (23-29nt). piRNAs protect the zygotic genome against infection by retroviruses that can come from the surrounding follicle cells or endogenous transposons within the female germline(4). piRNAs function together with the well-characterized SMAUG to direct the CCR4 deadenylase to specific mRNAs, thus facilitating maternal mRNA deadenylation and decay(7). The cytoplasmic localization of piRNAs in early development is consistent with their role in deadenylation of maternal mRNAs. The exact role of endogenous siRNAs in early development is not well-defined. Based on the observation that the mRNA profiles of wild type and *Dgcr8* null mouse oocytes were identical, it was proposed that endo-siRNAs, rather than miRNAs, are responsible for the *Dicer* knockout phenotype observed in mice(53). Thus, endo-siRNAs and piRNAs are primarily involved in gametogenesis and very early development whereas miRNAs appear to get involved in later developmental stages(54,55). Our data is consistent with this view in that transposon-derived transcripts are more abundant in 0-1h embryos (both fractionated and unfractionated, Figure 1B) whereas miRNA expression is up-regulated in 7-8h embryos. Endogenous siRNAs are well-known for their role in heterochromatin formation in the nucleus(56). The identification of endo-siRNAs in the cytoplasm points to two possibilities (1) the process of maturation as they are produced from dsRNAs in cytoplasm; (2) a potential role for these small RNAs in post-transcriptional gene regulation.

Transposon-derived siRNAs and piRNAs differ from each other in two ways: (i) 21-nt transposon-derived siRNAs are highly expressed in 7-8h embryos whereas piRNAs abound in 0-1h embryos (Figure 2–3), and (ii) siRNAs are mainly associated with the complexes in the mRNP fraction while piRNAs are largely associated with the polysomal complexes. It is quite interesting that both siRNAs and piRNAs are produced from the same transposons, yet they interact with the cellular translational machinery quite differently Another interesting point is that we detected a strand-bias in the reads obtained from the dual-strand transposon 42AB in 0-1h embryos especially in the polysome fraction (Figure 5B). It remains to be investigated whether there is a functional relationship between the biased-production of piRNAs and development.

We previously reported that tRNAs also serve as templates for small RNAs,.e., ~28-nt tRNA-derived fragments (tRFs)(35). These small RNAs are expressed at both stages (Figure 1B), and in fact throughout the embryonic development and in mature flies as well. They are mainly associated with non-polysomal fractions, resembling transposon-derived siRNAs. Studies on tRF function mainly focused on miRNA-like regulatory functions. In *Drosophila*, tRFs are immunoprecipitated with anti-AGO1 antibody(57). Interestingly, they inhibit cap-dependent translation initiation(58). It is unknown however whether tRFs are specifically involved in modulation of translation at the MZT stage.

## ACKNOWLEDGEMENTS

We thank Prof. Dr. C.-P. David Tu of Pennsylvania State University for providing the *Drosophila* P2 strain. We also thank the İYTE Biotechnology Research Center for the instrumental support.

---

**Supporting Figure 1.**
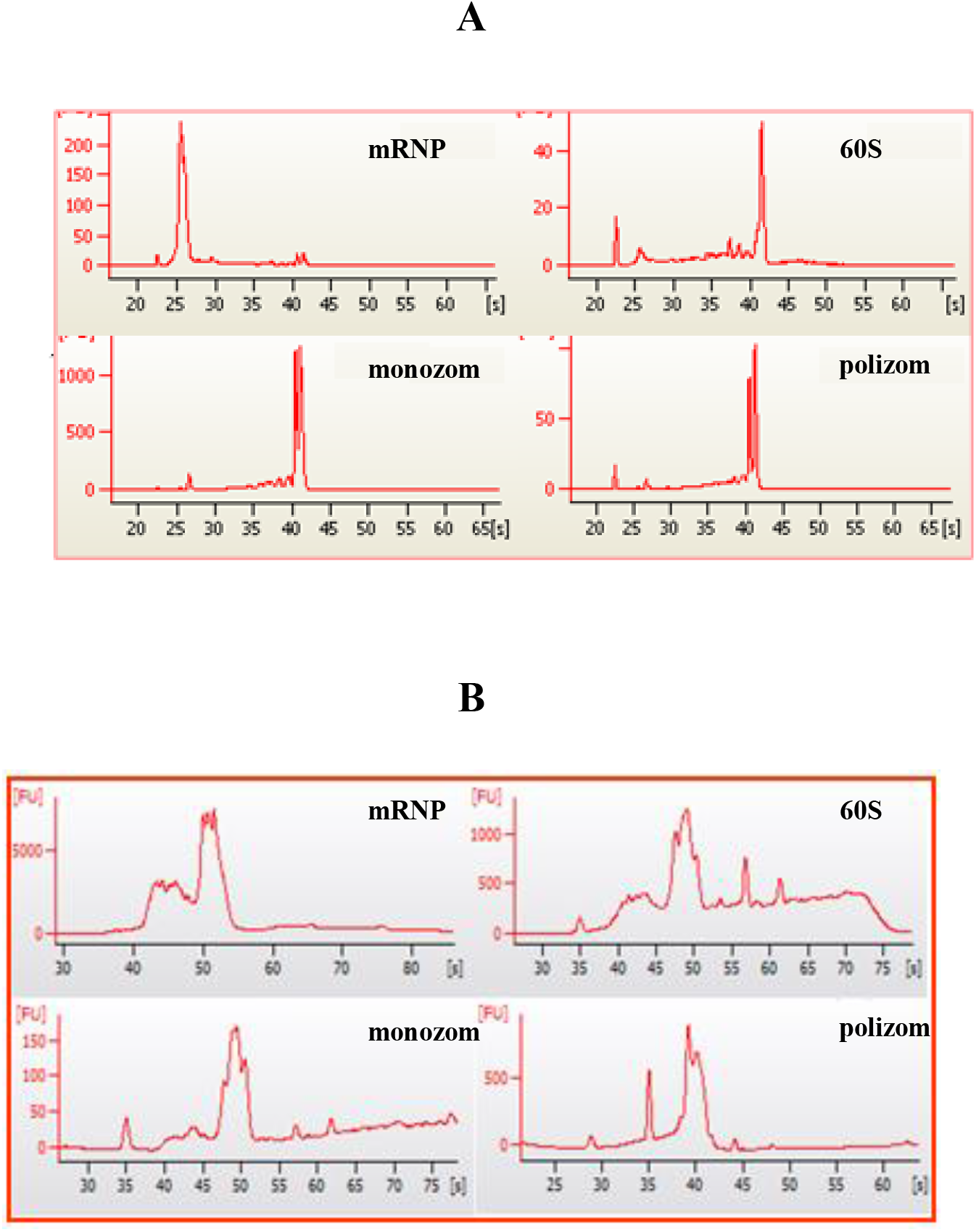
Total RNAs phenol extracted from fractionated embryos were run on Bionalyzer to assess the quality and size distribution of RNAs. Electropherograms of total RNAs using Agilent RNA 6000 Nano kit (**A**) and small RNA kit (**B**).

**Supporting Figure 2.**
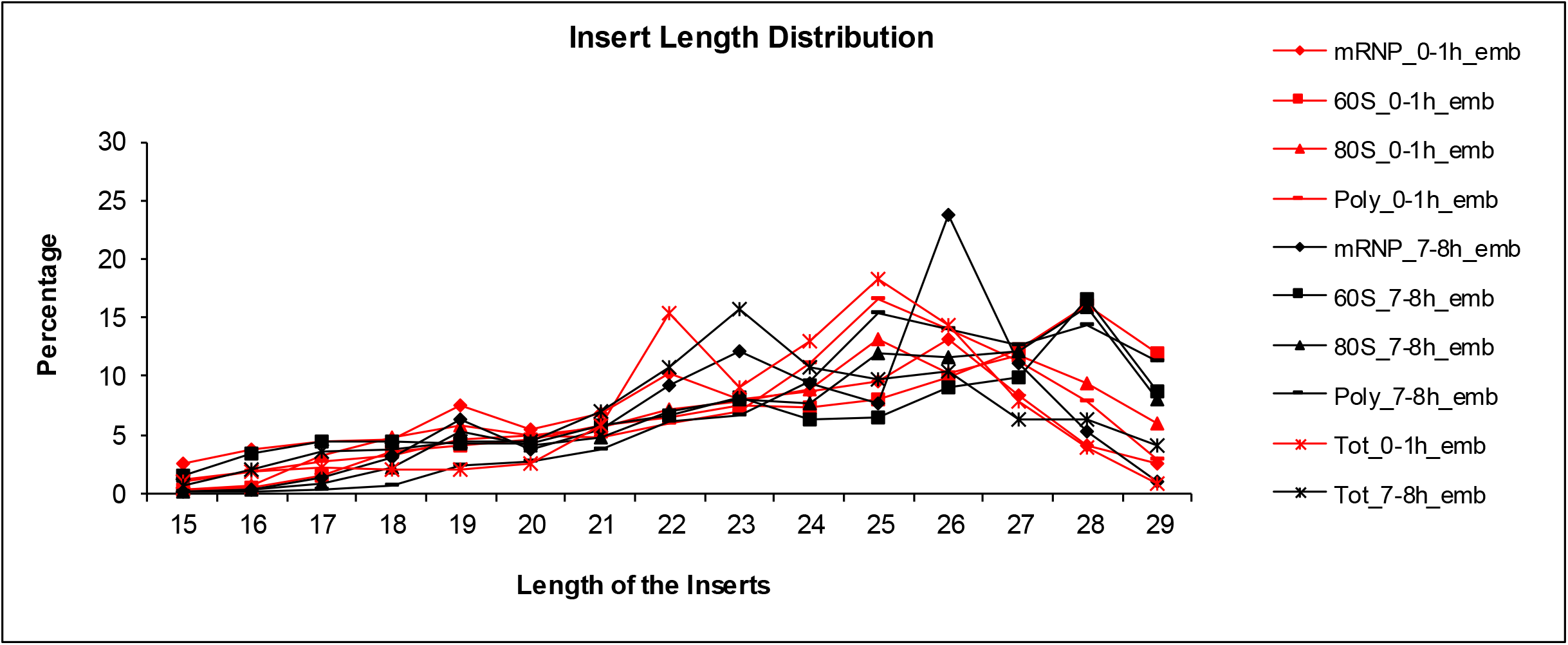

**Supporting Table 1.**
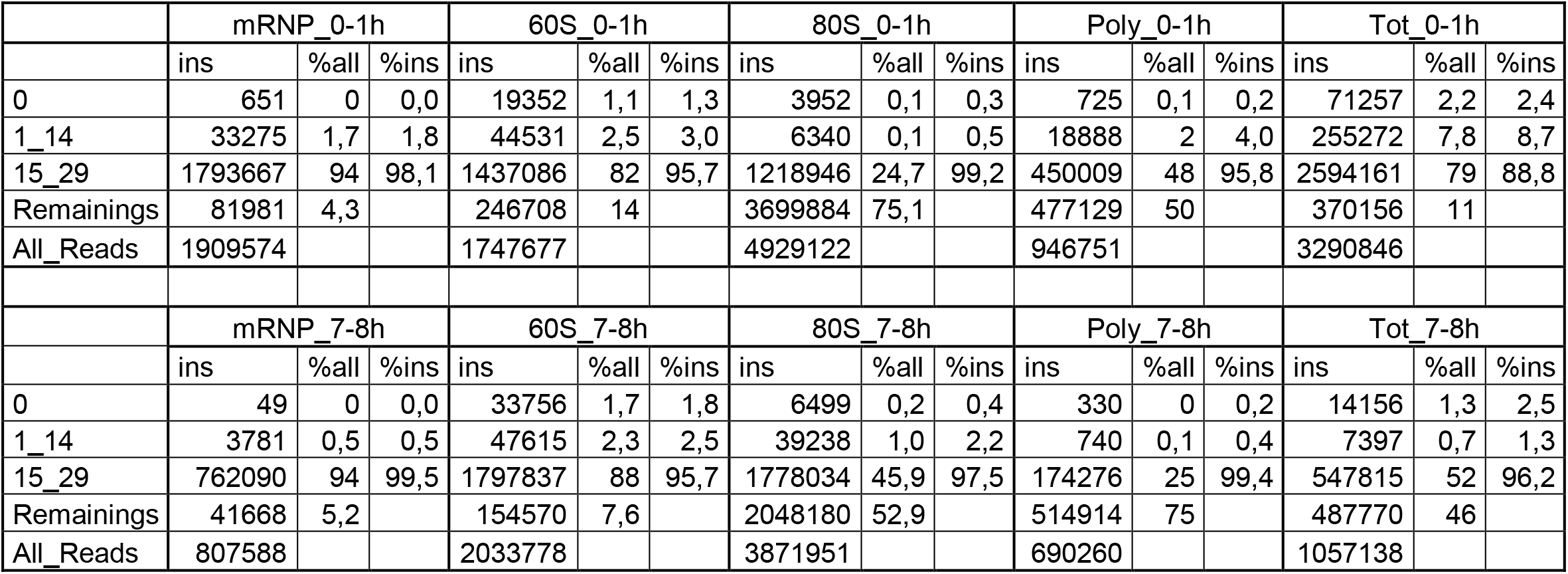
The percentage of hairpin and mature miRNA mapped reads in all reads mapped to miRNAs. For each fraction the percentage of each hairpin-mapped and mature miRNA reads calculated in total reads mapped to hairpin sequences.

**Coşacak et al Supporting Table 1**
|  | mRNP_0-1h |  |  | 60S_0-1h |  |  | 80S_0-1h |  |  | Poly_0-1h |  |  | Tot_0-1h |  |  |
| --- | --- | --- | --- | --- | --- | --- | --- | --- | --- | --- | --- | --- | --- | --- | --- |
|  | ins | %all | %ins | ins | %all | %ins | ins | %all | %ins | ins | %all | %ins | ins | %all | %ins |
| 0 | 651 | 0 | 0,0 | 19352 | 1,1 | 1,3 | 3952 | 0,1 | 0,3 | 725 | 0,1 | 0,2 | 71257 | 2,2 | 2,4 |
| 1_14 | 33275 | 1,7 | 1,8 | 44531 | 2,5 | 3,0 | 6340 | 0,1 | 0,5 | 18888 | 2 | 4,0 | 255272 | 7,8 | 8,7 |
| 15_29 | 1793667 | 94 | 98,1 | 1437086 | 82 | 95,7 | 1218946 | 24,7 | 99,2 | 450009 | 48 | 95,8 | 2594161 | 79 | 88,8 |
| Remainings | 81981 | 4,3 |  | 246708 | 14 |  | 3699884 | 75,1 |  | 477129 | 50 |  | 370156 | 11 |  |
| All_Reads | 1909574 |  |  | 1747677 |  |  | 4929122 |  |  | 946751 |  |  | 3290846 |  |  |
|  | mRNP_7-8h |  |  | 60S_7-8h |  |  | 80S_7-8h |  |  | Poly_7-8h |  |  | Tot_7-8h |  |  |
|  | ins | %all | %ins | ins | %all | %ins | ins | %all | %ins | ins | %all | %ins | ins | %all | %ins |
| 0 | 49 | 0 | 0,0 | 33756 | 1,7 | 1,8 | 6499 | 0,2 | 0,4 | 330 | 0 | 0,2 | 14156 | 1,3 | 2,5 |
| 1_14 | 3781 | 0,5 | 0,5 | 47615 | 2,3 | 2,5 | 39238 | 1,0 | 2,2 | 740 | 0,1 | 0,4 | 7397 | 0,7 | 1,3 |
| 15_29 | 762090 | 94 | 99,5 | 1797837 | 88 | 95,7 | 1778034 | 45,9 | 97,5 | 174276 | 25 | 99,4 | 547815 | 52 | 96,2 |
| Remainings | 41668 | 5,2 |  | 154570 | 7,6 |  | 2048180 | 52,9 |  | 514914 | 75 |  | 487770 | 46 |  |
| All_Reads | 807588 |  |  | 2033778 |  |  | 3871951 |  |  | 690260 |  |  | 1057138 |  |  |

## REFERENCES

1. De Renzis, S., Elemento, O., Tavazoie, S. and Wieschaus, E.F. (2007) Unmasking activation of the zygotic genome using chromosomal deletions in the Drosophila embryo. PLoS Biol, 5, e117.

2. Tadros, W., Goldman, A.L., Babak, T., Menzies, F., Vardy, L., Orr-Weaver, T., Hughes, T.R., Westwood, J.T., Smibert, C.A. and Lipshitz, H.D. (2007) SMAUG is a major regulator of maternal mRNA destabilization in Drosophila and its translation is activated by the PAN GU kinase. Dev Cell, 12, 143–155.

3. Alberti, C. and Cochella, L. (2017) A framework for understanding the roles of miRNAs in animal development. Development, 144, 2548–2559.

4. Bourc’his, D. and Voinnet, O. (2010) A small-RNA perspective on gametogenesis, fertilization, and early zygotic development. Science, 330, 617–622.

5. Bushati, N., Stark, A., Brennecke, J. and Cohen, S.M. (2008) Temporal reciprocity of miRNAs and their targets during the maternal-to-zygotic transition in Drosophila. Curr Biol, 18, 501–506.

6. Ma, J., Flemr, M., Stein, P., Berninger, P., Malik, R., Zavolan, M., Svoboda, P. and Schultz, R.M. (2010) MicroRNA activity is suppressed in mouse oocytes. Curr Biol, 20, 265–270.

7. Rouget, C., Papin, C., Boureux, A., Meunier, A.C., Franco, B., Robine, N., Lai, E.C., Pelisson, A. and Simonelig, M. (2010) Maternal mRNA deadenylation and decay by the piRNA pathway in the early Drosophila embryo. Nature, 467, 1128–1132.

8. Carthew, R.W. and Sontheimer, E.J. (2009) Origins and Mechanisms of miRNAs and siRNAs. Cell, 136, 642–655.

9. Ghildiyal, M. and Zamore, P.D. (2009) Small silencing RNAs: an expanding universe. Nat Rev Genet, 10, 94–108.

10. Weick, E.M. and Miska, E.A. (2014) piRNAs: from biogenesis to function. Development, 141, 3458–3471.

11. Chendrimada, T.P., Finn, K.J., Ji, X., Baillat, D., Gregory, R.I., Liebhaber, S.A., Pasquinelli, A.E. and Shiekhattar, R. (2007) MicroRNA silencing through RISC recruitment of eIF6. Nature, 447, 823–828.

12. Guo, H., Ingolia, N.T., Weissman, J.S. and Bartel, D.P. (2010) Mammalian microRNAs predominantly act to decrease target mRNA levels. Nature, 466, 835–840.

13. Kiriakidou, M., Tan, G.S., Lamprinaki, S., De Planell-Saguer, M., Nelson, P.T. and Mourelatos, Z. (2007) An mRNA m7G cap binding-like motif within human Ago2 represses translation. Cell, 129, 1141–1151.

14. Petersen, C.P., Bordeleau, M.E., Pelletier, J. and Sharp, P.A. (2006) Short RNAs repress translation after initiation in mammalian cells. Mol Cell, 21, 533–542.

15. Aravin, A.A., Lagos-Quintana, M., Yalcin, A., Zavolan, M., Marks, D., Snyder, B., Gaasterland, T., Meyer, J. and Tuschl, T. (2003) The small RNA profile during Drosophila melanogaster development. Dev Cell, 5, 337–350.

16. Okamura, K. and Lai, E.C. (2008) Endogenous small interfering RNAs in animals. Nat Rev Mol Cell Biol, 9, 673–678.

17. Barckmann, B., Pierson, S., Dufourt, J., Papin, C., Armenise, C., Port, F., Grentzinger, T., Chambeyron, S., Baronian, G., Desvignes, J.P. et al. (2015) Aubergine iCLIP Reveals piRNA-Dependent Decay of mRNAs Involved in Germ Cell Development in the Early Embryo. Cell Rep, 12, 1205–1216.

18. Claycomb, J.M. (2014) Ancient endo-siRNA pathways reveal new tricks. Curr Biol, 24, R703–715.

19. Yuan, S., Schuster, A., Tang, C., Yu, T., Ortogero, N., Bao, J., Zheng, H. and Yan, W. (2016) Sperm-borne miRNAs and endo-siRNAs are important for fertilization and preimplantation embryonic development. Development, 143, 635–647.

20. Zhang, H., Liu, J., Tai, Y., Zhang, X., Zhang, J., Liu, S., Lv, J., Liu, Z. and Kong, Q. (2017) Identification and characterization of L1-specific endo-siRNAs essential for early embryonic development in pig. Oncotarget, 8, 23167–23176.

21. Aboobaker, A.A., Tomancak, P., Patel, N., Rubin, G.M. and Lai, E.C. (2005) Drosophila microRNAs exhibit diverse spatial expression patterns during embryonic development. Proc Natl Acad Sci U S A, 102, 18017–18022.

22. Leaman, D., Chen, P.Y., Fak, J., Yalcin, A., Pearce, M., Unnerstall, U., Marks, D.S., Sander, C., Tuschl, T. and Gaul, U. (2005) Antisense-mediated depletion reveals essential and specific functions of microRNAs in Drosophila development. Cell, 121, 1097–1108.

23. Chen, Y.W., Song, S., Weng, R., Verma, P., Kugler, J.M., Buescher, M., Rouam, S. and Cohen, S.M. (2014) Systematic study of Drosophila microRNA functions using a collection of targeted knockout mutations. Dev Cell, 31, 784–800.

24. Tang, F., Kaneda, M., O’Carroll, D., Hajkova, P., Barton, S.C., Sun, Y.A., Lee, C., Tarakhovsky, A., Lao, K. and Surani, M.A. (2007) Maternal microRNAs are essential for mouse zygotic development. Genes Dev, 21, 644–648.

25. Soni, K., Choudhary, A., Patowary, A., Singh, A.R., Bhatia, S., Sivasubbu, S., Chandrasekaran, S. and Pillai, B. (2013) miR-34 is maternally inherited in Drosophila melanogaster and Danio rerio. Nucleic Acids Res, 41, 4470–4480.

26. Svoboda, P. and Flemr, M. (2010) The role of miRNAs and endogenous siRNAs in maternal-to-zygotic reprogramming and the establishment of pluripotency. EMBO Rep, 11, 590–597.

27. Fu, S., Nien, C.Y., Liang, H.L. and Rushlow, C. (2014) Co-activation of microRNAs by Zelda is essential for early Drosophila development. Development, 141, 2108–2118.

28. Lee, M., Choi, Y., Kim, K., Jin, H., Lim, J., Nguyen, T.A., Yang, J., Jeong, M., Giraldez, A.J., Yang, H. et al. (2014) Adenylation of maternally inherited microRNAs by Wispy. Mol Cell, 56, 696–707.

29. Ha, M. and Kim, V.N. (2014) Regulation of microRNA biogenesis. Nat Rev Mol Cell Biol, 15, 509–524.

30. Watanabe, T. and Lin, H. (2014) Posttranscriptional regulation of gene expression by Piwi proteins and piRNAs. Mol Cell, 56, 18–27.

31. Liu, J., Valencia-Sanchez, M.A., Hannon, G.J. and Parker, R. (2005) MicroRNA-dependent localization of targeted mRNAs to mammalian P-bodies. Nat Cell Biol, 7, 719–723.

32. Maroney, P.A., Yu, Y., Fisher, J. and Nilsen, T.W. (2006) Evidence that microRNAs are associated with translating messenger RNAs in human cells. Nat Struct Mol Biol, 13, 1102–1107.

33. Molotski, N. and Soen, Y. (2012) Differential association of microRNAs with polysomes reflects distinct strengths of interactions with their mRNA targets. RNA, 18, 1612–1623.

34. Wu, P.H., Isaji, M. and Carthew, R.W. (2013) Functionally diverse microRNA effector complexes are regulated by extracellular signaling. Mol Cell, 52, 113–123.

35. Goktas, C., Yigit, H., Cosacak, M.I. and Akgul, B. (2017) Differentially Expressed tRNA-Derived Small RNAs Co-Sediment Primarily with Non-Polysomal Fractions in Drosophila. Genes (Basel), 8.

36. de Hoon, M.J., Taft, R.J., Hashimoto, T., Kanamori-Katayama, M., Kawaji, H., Kawano, M., Kishima, M., Lassmann, T., Faulkner, G.J., Mattick, J.S. et al. (2010) Cross-mapping and the identification of editing sites in mature microRNAs in high-throughput sequencing libraries. Genome Res, 20, 257–264.

37. Jurka, J., Kapitonov, V.V., Pavlicek, A., Klonowski, P., Kohany, O. and Walichiewicz, J. (2005) Repbase Update, a database of eukaryotic repetitive elements. Cytogenet Genome Res, 110, 462–467.

38. Brennecke, J., Aravin, A.A., Stark, A., Dus, M., Kellis, M., Sachidanandam, R. and Hannon, G.J. (2007) Discrete small RNA-generating loci as master regulators of transposon activity in Drosophila. Cell, 128, 1089–1103.

39. de Hoon, M.J., Imoto, S., Nolan, J. and Miyano, S. (2004) Open source clustering software. Bioinformatics, 20, 1453–1454.

40. Saldanha, A.J. (2004) Java Treeview--extensible visualization of microarray data. Bioinformatics, 20, 3246–3248.

41. Altuvia, Y., Landgraf, P., Lithwick, G., Elefant, N., Pfeffer, S., Aravin, A., Brownstein, M.J., Tuschl, T. and Margalit, H. (2005) Clustering and conservation patterns of human microRNAs. Nucleic Acids Res, 33, 2697–2706.

42. Baskerville, S. and Bartel, D.P. (2005) Microarray profiling of microRNAs reveals frequent coexpression with neighboring miRNAs and host genes. RNA, 11, 241–247.

43. Wang, J., Haubrock, M., Cao, K.M., Hua, X., Zhang, C.Y., Wingender, E. and Li, J. (2011) Regulatory coordination of clustered microRNAs based on microRNA-transcription factor regulatory network. BMC Syst Biol, 5, 199.

44. Kawahara, Y., Zinshteyn, B., Sethupathy, P., Iizasa, H., Hatzigeorgiou, A.G. and Nishikura, K. (2007) Redirection of silencing targets by adenosine-to-inosine editing of miRNAs. Science, 315, 1137–1140.

45. Grivna, S.T., Pyhtila, B. and Lin, H. (2006) MIWI associates with translational machinery and PIWI-interacting RNAs (piRNAs) in regulating spermatogenesis. Proc Natl Acad Sci U S A, 103, 13415–13420.

46. Ghildiyal, M., Seitz, H., Horwich, M.D., Li, C., Du, T., Lee, S., Xu, J., Kittler, E.L., Zapp, M.L., Weng, Z. et al. (2008) Endogenous siRNAs derived from transposons and mRNAs in Drosophila somatic cells. Science, 320, 1077–1081.

47. Aravin, A., Gaidatzis, D., Pfeffer, S., Lagos-Quintana, M., Landgraf, P., Iovino, N., Morris, P., Brownstein, M.J., Kuramochi-Miyagawa, S., Nakano, T. et al. (2006) A novel class of small RNAs bind to MILI protein in mouse testes. Nature, 442, 203–207.

48. Klattenhoff, C., Xi, H., Li, C., Lee, S., Xu, J., Khurana, J.S., Zhang, F., Schultz, N., Koppetsch, B.S., Nowosielska, A. et al. (2009) The Drosophila HP1 homolog Rhino is required for transposon silencing and piRNA production by dual-strand clusters. Cell, 138, 1137–1149.

49. Chung, W.J., Okamura, K., Martin, R. and Lai, E.C. (2008) Endogenous RNA interference provides a somatic defense against Drosophila transposons. Curr Biol, 18, 795–802.

50. Giraldez, A.J., Mishima, Y., Rihel, J., Grocock, R.J., Van Dongen, S., Inoue, K., Enright, A.J. and Schier, A.F. (2006) Zebrafish MiR-430 promotes deadenylation and clearance of maternal mRNAs. Science, 312, 75–79.

51. Tchurikov, N.A. and Kretova, O.V. (2011) Both piRNA and siRNA pathways are silencing transcripts of the suffix element in the Drosophila melanogaster germline and somatic cells. PLoS One, 6, e21882.

52. Zong, Q., Schummer, M., Hood, L. and Morris, D.R. (1999) Messenger RNA translation state: the second dimension of high-throughput expression screening. Proc Natl Acad Sci U S A, 96, 10632–10636.

53. Suh, N., Baehner, L., Moltzahn, F., Melton, C., Shenoy, A., Chen, J. and Blelloch, R. (2010) MicroRNA function is globally suppressed in mouse oocytes and early embryos. Curr Biol, 20, 271–277.

54. Suh, N. and Blelloch, R. (2011) Small RNAs in early mammalian development: from gametes to gastrulation. Development, 138, 1653–1661.

55. Dallaire, A. and Simard, M.J. (2016) The implication of microRNAs and endo-siRNAs in animal germline and early development. Dev Biol, 416, 18–25.

56. Fagegaltier, D., Bouge, A.L., Berry, B., Poisot, E., Sismeiro, O., Coppee, J.Y., Theodore, L., Voinnet, O. and Antoniewski, C. (2009) The endogenous siRNA pathway is involved in heterochromatin formation in Drosophila. Proc Natl Acad Sci U S A, 106, 21258–21263.

57. Karaiskos, S., Naqvi, A.S., Swanson, K.E. and Grigoriev, A. (2015) Age-driven modulation of tRNA-derived fragments in Drosophila and their potential targets. Biol Direct, 10, 51.

58. Sobala, A. and Hutvagner, G. (2013) Small RNAs derived from the 5′ end of tRNA can inhibit protein translation in human cells. RNA Biol, 10, 553–563.

